# PncA from bacteria improves diet-induced NAFLD by enabling the transition from NAM to NA in mice

**DOI:** 10.1101/2021.12.04.471202

**Authors:** Shengyu Feng, Liuling Guo, Hailiang Liu

**Affiliations:** Institute for Regenerative Medicine, Shanghai East Hospital, Tongji University School of Medicine, Shanghai, 200123, China; Key Laboratory of Xinjiang Phytomedicine Resource and Utilization of Ministry of Education, College of Life Sciences, Shihezi University, Shihezi, 832003, China

## Abstract

Nicotinamide adenine dinucleotide (NAD^+^) is crucial for energy metabolism, oxidative stress, DNA damage repair, longevity regulation, and several signaling processes. To date, three NAD^+^ synthesis pathways have been found in microbiota and hosts, but the potential relationship between gut microbiota and their hosts in regulating NAD^+^ homeostasis remains unknown. Here, we show that an analog of the first-line tuberculosis drug pyrazinamide (a bacterial NAD^+^ synthesis inhibitor) affected NAD^+^ levels in the intestines and liver of mice and disrupted the intestinal microecological balance. Furthermore, using microbiota expressing the pyrazinamidase/nicotinamidase (*PncA*) gene, which is a target of pyrazinamide, hepatic NAD^+^ levels were greatly increased and significantly increased compared with other NAD^+^ precursors, and diet-induced non-alcoholic fatty liver disease (NAFLD) in mice was improved. Overall, the *PncA* gene in microbiota plays an important role in regulating NAD^+^ synthesis in the host, thereby providing a potential target for modulating the host’s NAD^+^ level.

**Highlights:**

1. PncA inhibitors disrupt gut microbiome homeostasis and reduce host NAD^+^ levels but do not affect NAD^+^ levels in cultured cells
2. *PncA* gene in microbiota affects host liver NAD metabolism
3. PncA affects lipid metabolism-related genes and metabolites in mice with NAFLD
4. Diet-induced NAFLD is improved by PncA overexpression in the liver of mice

**Graphical abstract:** 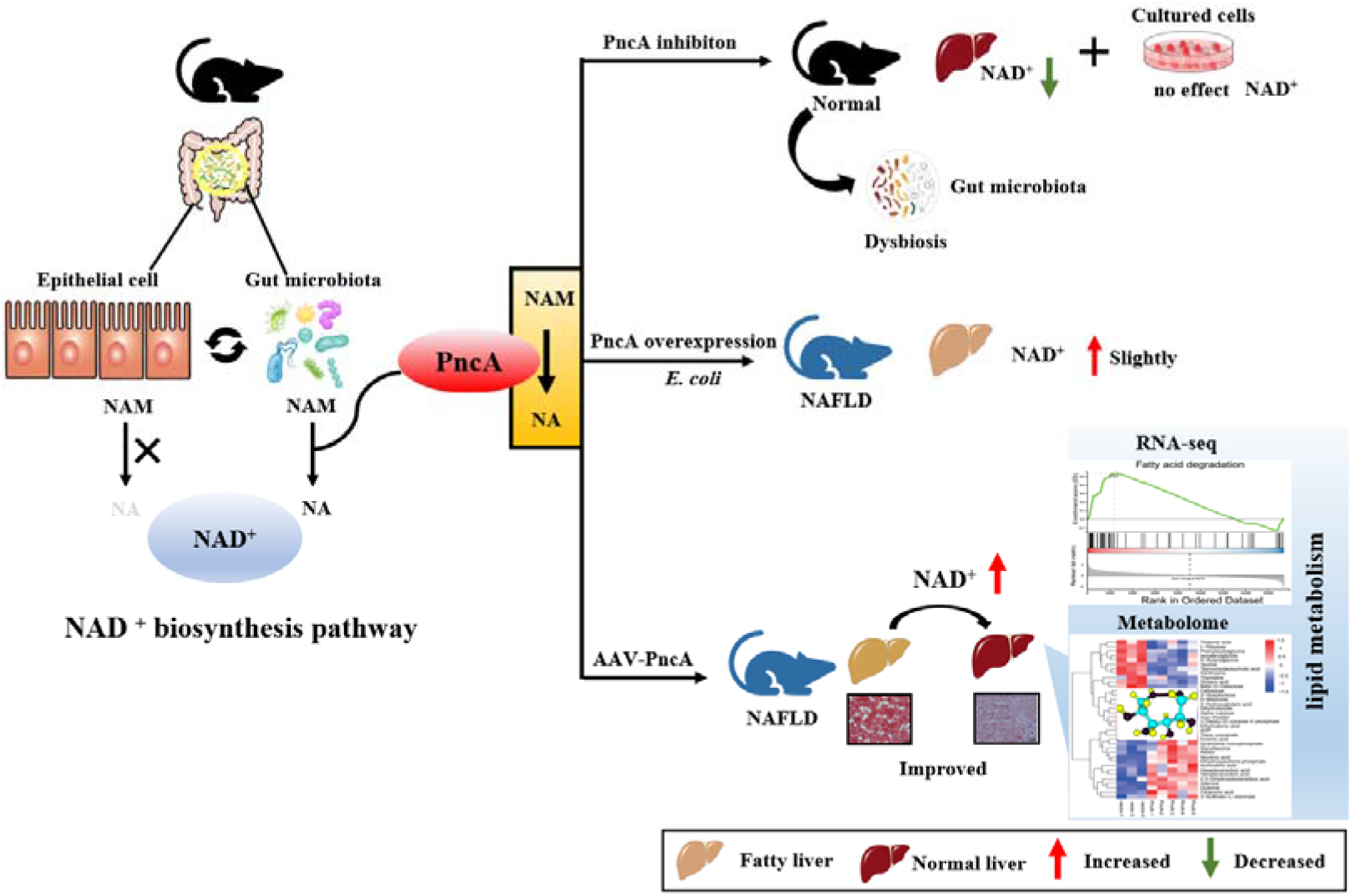

## Main

As early as 110 years ago, NAD^+^ was discovered by the British biochemists Arthur Harden and William John Young as a “coenzyme” involved in yeast-mediated alcohol fermentation ^1^. Over the next ∼30 years, the chemical composition of this “coenzyme” was determined, and it was found to participate in several redox reactions together with NADH. Although the role of NAD^+^ in oxidation-reduction reactions is well understood, it was not until the last 10 years that the function of NAD was completely elucidated. The sirtuin family (SIRTs), poly (ADP-ribose) polymerases (PARPs), and cyclic ADP-ribose syntheses (cADPRSs) are NAD^+^-dependent enzymes, and NAD^+^ regulates downstream metabolic pathways by influencing the activity of these enzymes and acting as a metabolic sensor in cells ^2-5^. In addition, these NAD^+^-dependent proteins play an important role in a variety of biological processes, such as metabolism, signal transduction, oxidative stress, cognitive decline, and other aging-related physiological processes. In recent years, important studies have revealed that NAD^+^ binds to the 5’ end of mRNA, thereby regulating the transcription initiation of genes. However, there is still no definitive conclusion on how this process is regulated or its significance for organisms ^6, 7^.

NAD^+^ is a small molecule required by almost all organisms and is one of the most abundant molecules in the human body, participating in more than 500 different enzymatic reactions. The NAD^+^ amount in an adult is approximately 3 g. We summarized the NAD^+^ synthesis and consumption pathways in mammals and human intestinal flora (Fig. 1). In mammals, there are three main synthesis pathways for NAD^+^: the de novo synthesis pathway using tryptophan, the “salvage” pathway using nicotinamide (NAM), and the “Preiss–Handler” pathway using nicotinic acid (NA). Because there are several mechanisms to synthesize NAD^+^ in mammals, the specific pathways used for different cell types and the most effective pathway remain unclear ^8, 9^. In the intestinal flora, there is an additional important pathway that converts NAM into NA, combining the “salvage” and “Preiss–Handler” pathways. This is known as deamidation and is catalyzed by pyrazinamidase/nicotinamidase (PncA)^10^. This process has recently been shown to be an important step for the intestinal flora to regulate the host’s NAD^+^ level ^11^, and this pathway performed by PncA is considered to have been abandoned during biological evolution because no homolog of the gene has been identified in higher organisms. However, it is also possible that this gene was mutated during the evolution of higher organisms, resulting in its low genetic similarity with lower organisms. As a result, no enzyme catalyzing the conversion of NAM to NA has been found in mammals to date ^9^. However, the genes involved in this process are highly conserved in different bacterial phyla (Supplementary Fig. S1), indicating their importance in the bacterial synthesis of NAD^+^. Based on these characteristics, pyrazinamide targeting the *PncA* gene was developed to treat *Mycobacterium tuberculosis* ^12^. In addition, Supplementary Fig. S1 shows other genes related to NAD^+^ synthesis in different types of bacteria.

**Fig. 1:**
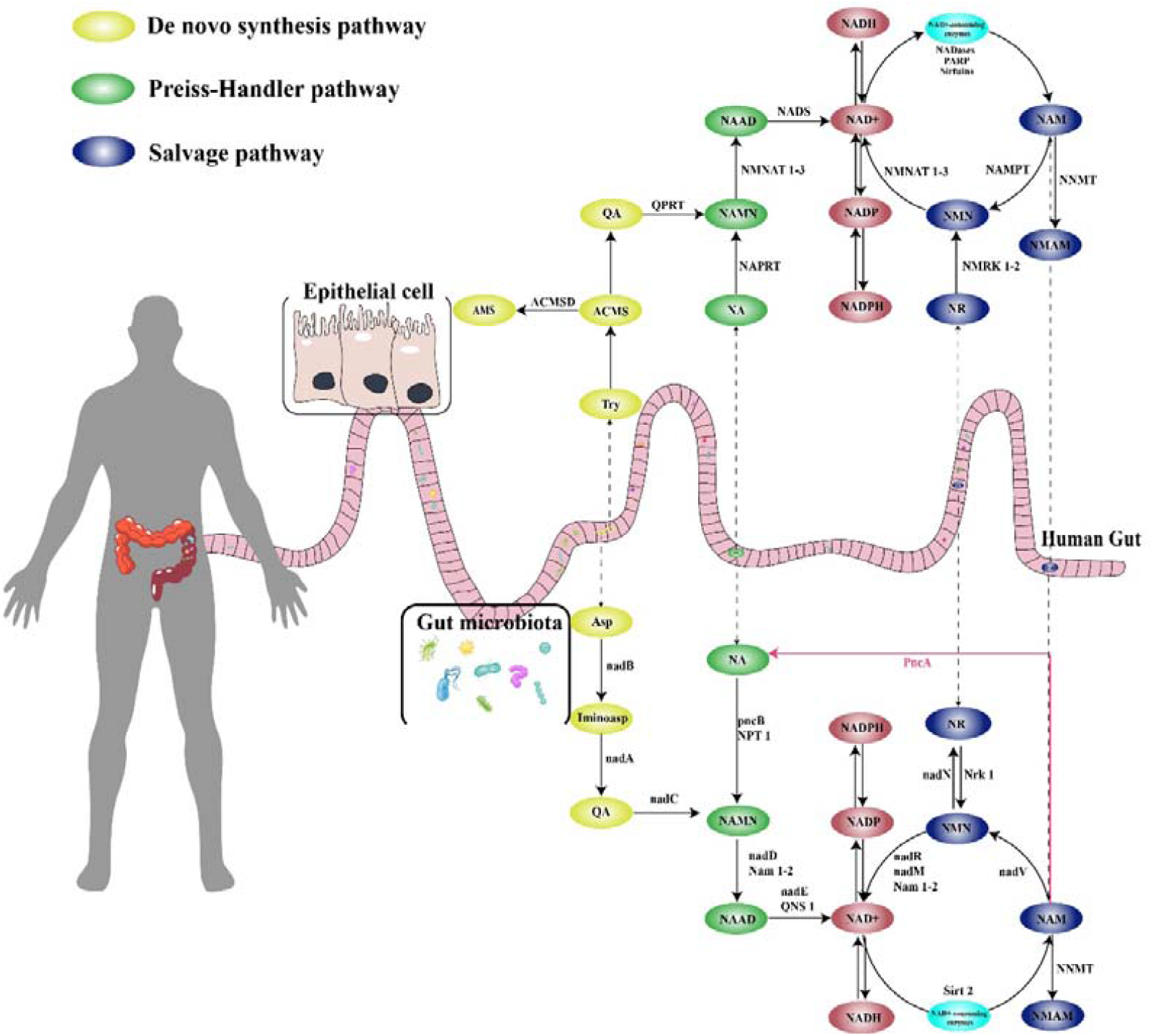
The synthesis and consumption pathway of NAD^+^ in human and intestinal flora. Two main pathways contribute to NAD synthesis in humans: the de novo synthesis and recovery of precursor synthesis pathways. The de novo synthesis pathway converts tryptophan to quinolinic acid (QA) through the kynurenine pathway and further synthesizes NAD^+^. The main precursors of NAD^+^ synthesis are NAM, NA, nicotinamide nucleoside, and NMN. NAD+-consuming enzymes mainly include sirtuins, PARPs, and CD38, and their substrates all contain NAM. In the intestinal flora, the de novo synthesis of NAD^+^ mainly uses aspartic acid rather than tryptophan, which is used in humans. Although the two organisms synthesize different substances de novo, they share the same precursors for NAD^+^ synthesis. Bacteria also mainly use NAM, NA, nicotinamide nucleoside, and NMN. Dependence on the same precursors must lead to competition or cooperation in NAD^+^ anabolism between humans and their gut flora. The synthetic pathway marked by the red line is specific to the gut flora and represents the deamidation pathway that converts NAM to NA.

Changes in liver metabolism are key factors contributing to the occurrence of liver diseases, among which non-alcoholic fatty liver disease (NAFLD) is the most common chronic liver disease and is closely related to metabolic syndrome. With substantial changes in diet and lifestyle, the prevalence of NAFLD has increased significantly worldwide ^13^. Owing to the complexity and heterogeneity of NAFLD, there are currently no effective FDA-approved drugs available for clinical use. Therefore, exploring promising therapeutic targets or strategies remains a priority ^14^. Several studies have shown that supplementation with the NAD^+^ precursors NAM nucleoside (NR) and nicotinamide mononucleotide (NMN) and ACMSD inhibition (an enzyme that blocks the de novo NAD^+^ synthesis pathway) significantly improve NAFLD in mice ^15-17^, suggesting NAD^+^ regulation in the diseased liver as a potential therapeutic target. During the development of NAFLD, the intestinal flora undergoes significant changes ^18^, and PncA in the intestinal flora plays an important role in NAD^+^ synthesis in mouse livers. Therefore, this paper aimed to further explore the potential significance of PncA in improving NAFLD in mice.

**Supplementary Figure 1:**
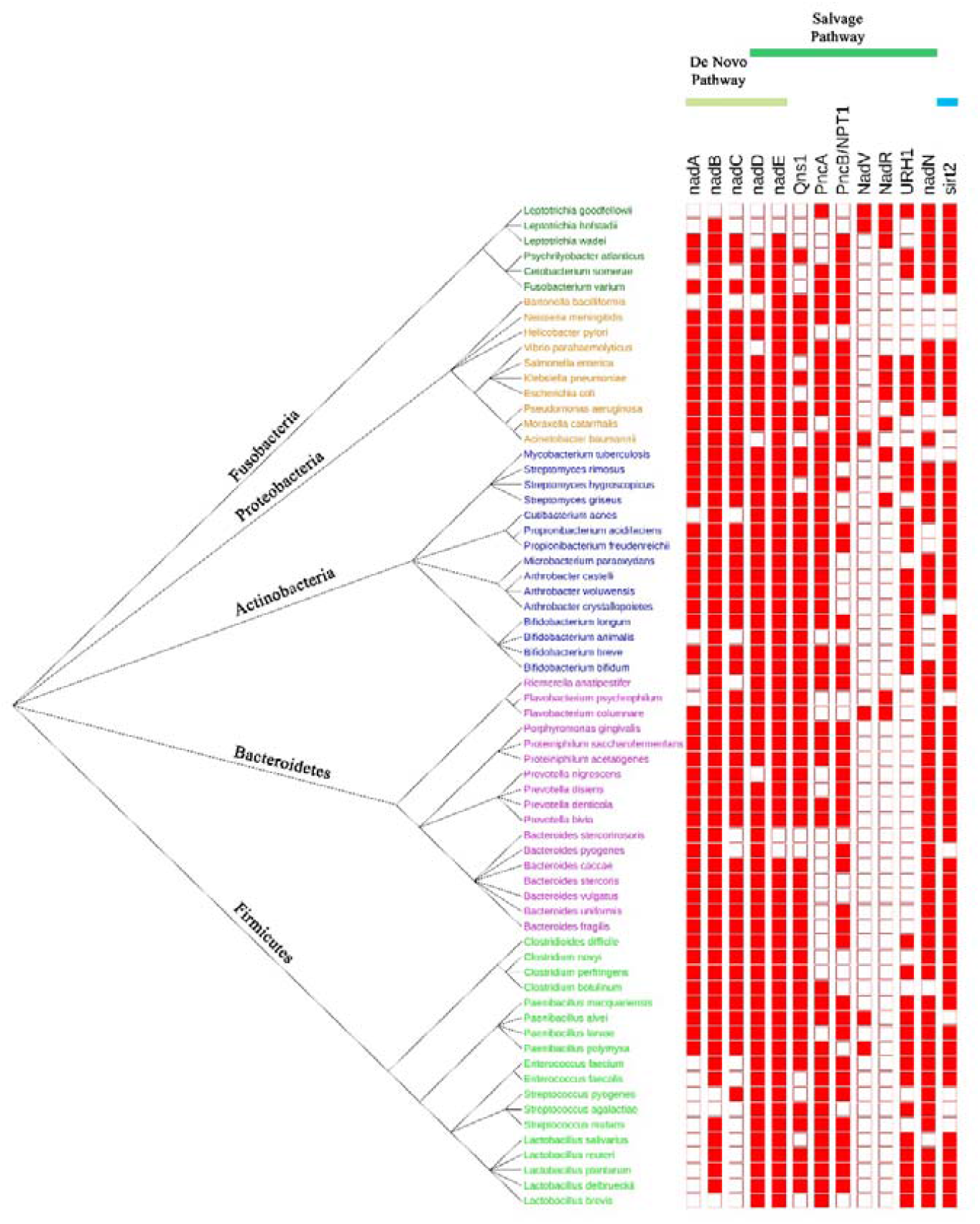
Genes associated with NAD^+^ synthesis in different taxa of bacteria. *nadA, nadB*, and *nadC* genes are involved in the NAD^+^ de novo synthesis pathway, and *nadD* and *nadE* genes are involved in both the de novo pathway and the NAD^+^ synthesis pathway using precursors. *PncA, PncB, nadR, nadC, nadR, URH1*, and *nadN* genes are only involved in the NAD^+^ synthesis pathway using precursors. The solid red box indicates that the species has the corresponding gene. Bacteria in different phylum groups are represented by different colors in the evolutionary tree.

## Results

### PncA inhibitors disrupt gut microbiome homeostasis and reduce host NAD^+^ levels

To verify whether the *PncA* gene in microbiota affects NAD^+^ synthesis in the host, we used the PncA inhibitor pyrazinecarbonitrile (PCN), which is an analog of the antibiotic pyrazinamide commonly used to treat tuberculosis. PCN has been reported to have a strong inhibitory effect on PncA enzyme activity ^19^. To verify the influence of PCN on bacterial growth *in vitro*, we selected several bacteria with different NAD^+^ synthesis pathways, including *Bifidobacterium longum* with de novo and deamidation synthesis pathways, *Akkermansia muciniphila* with the de novo synthesis pathway, and *Lactobacillus salivarius* with the deamidation synthesis pathway (Supplementary Fig. 1). These three types of bacteria are common flora in mammals. The growth rates of *Akkermansia muciniphila* and *Bifidobacterium longum* were mainly unaffected after 24 h of PCN treatment (Fig. 2A), whereas the growth of *Lactobacillus salivarius*, which relies only on the deamidation pathway, was strongly inhibited. This demonstrates that PCN has a strong inhibitory effect on deamidation pathway-dependent bacteria. Although *Bifidobacterium longum* expresses the *PncA* gene, PCN had a minimal effect on its growth, indicating that *Bifidobacterium longum* mainly synthesizes NAD^+^ through the de novo pathway.

**Fig. 2:**
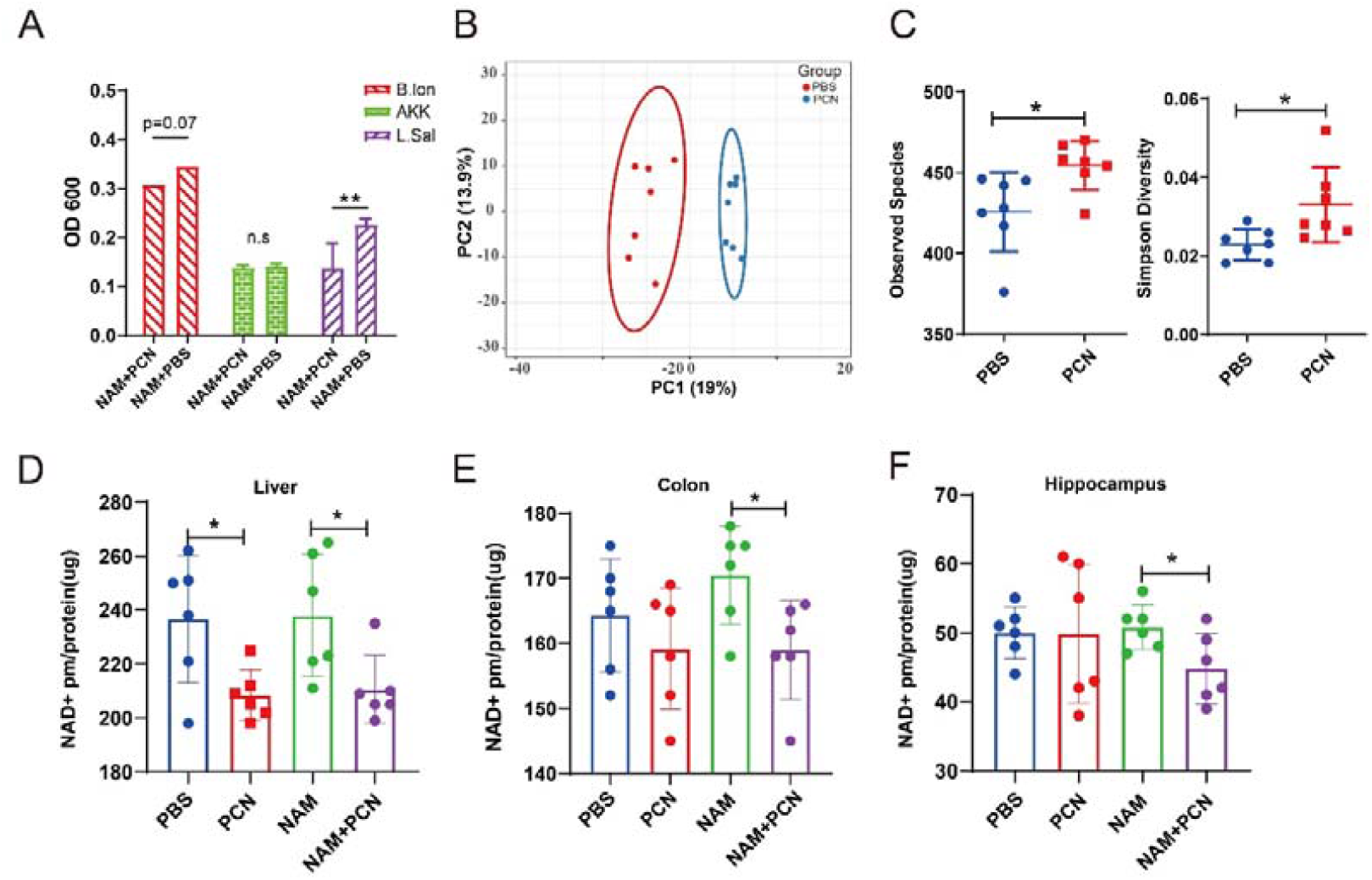
PncA inhibitors disrupt intestinal microbiome homeostasis and reduce host NAD+ levels. A. Effects of PCN on the growth of bacteria with different NAD^+^ synthesis pathways. B. PCA diagram of the intestinal microbe in mice treated with PCN. C. Intestinal microbial species and diversity after PCN treatment in mice. D–f. NAD^+^ level in the liver, intestine, and hippocampus of mice.

In the *in vivo* experiment, we treated mice with PCN and collected the feces of each group on the last day of the experiment. We then used sequencing technology to analyze the changes in the intestinal flora of mice. Principal component analysis (PCA) indicated that PCN treatment had a significant impact on the gut microbes in mice (Fig. 2B). Surprisingly, mice treated with PCN exhibited a significant increase in bacterial richness compared with control mice (Fig. 2C). As a potential explanation, we speculate that PCN disrupts the original balance in the intestinal flora, resulting in the massive expansion of some bacteria. After PCN treatment, the abundance of *Bacteroidales* increased significantly, whereas the abundance of *Clostridiales* decreased significantly (Supplementary Fig. 2A). At the species level, PCN significantly increased the abundance of *Helicobacter hepaticus, Clostridium cocleatum*, and *Bifidobacterium pseudolongum* (Supplementary Fig. 2B). Moreover, PCN significantly inhibited electron transfer, respiration, and other pathways closely related to NAD^+^ in the intestinal flora (Supplementary Fig. 2C). Together, these findings show that PCN has a significant impact on the intestinal flora.

**Supplementary Figure 2:**
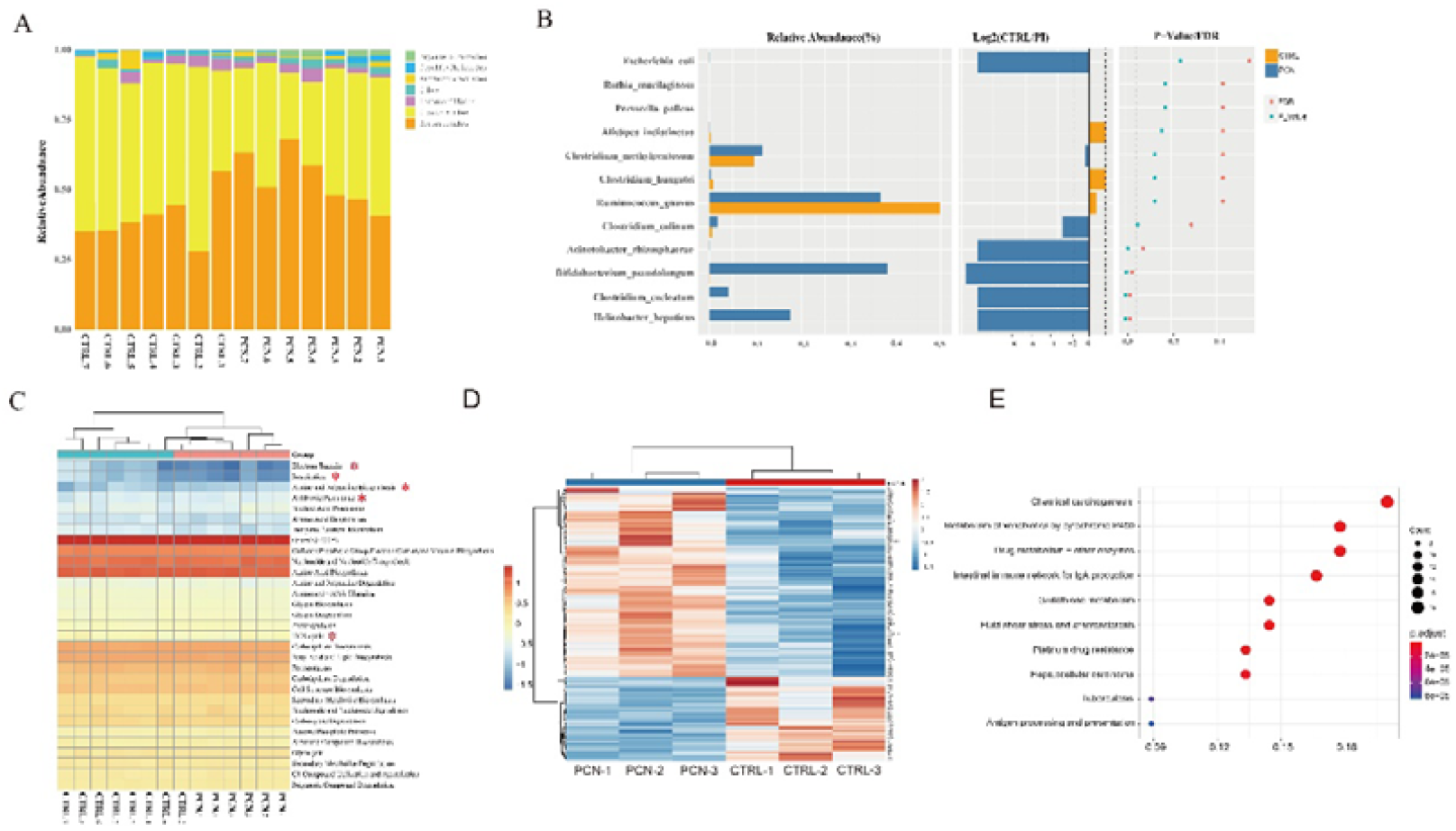
Effects of PCN on the intestinal microbiota and host. A. Distribution of bacteria at the order level. B. Distribution of bacteria at the species level. C. KEGG pathways enriched by bacteria. D. Heatmap of differentially expressed genes in the liver in the PCN group and control group. E. KEGG analysis of differently expressed genes.

**Supplementary Figure 3:**
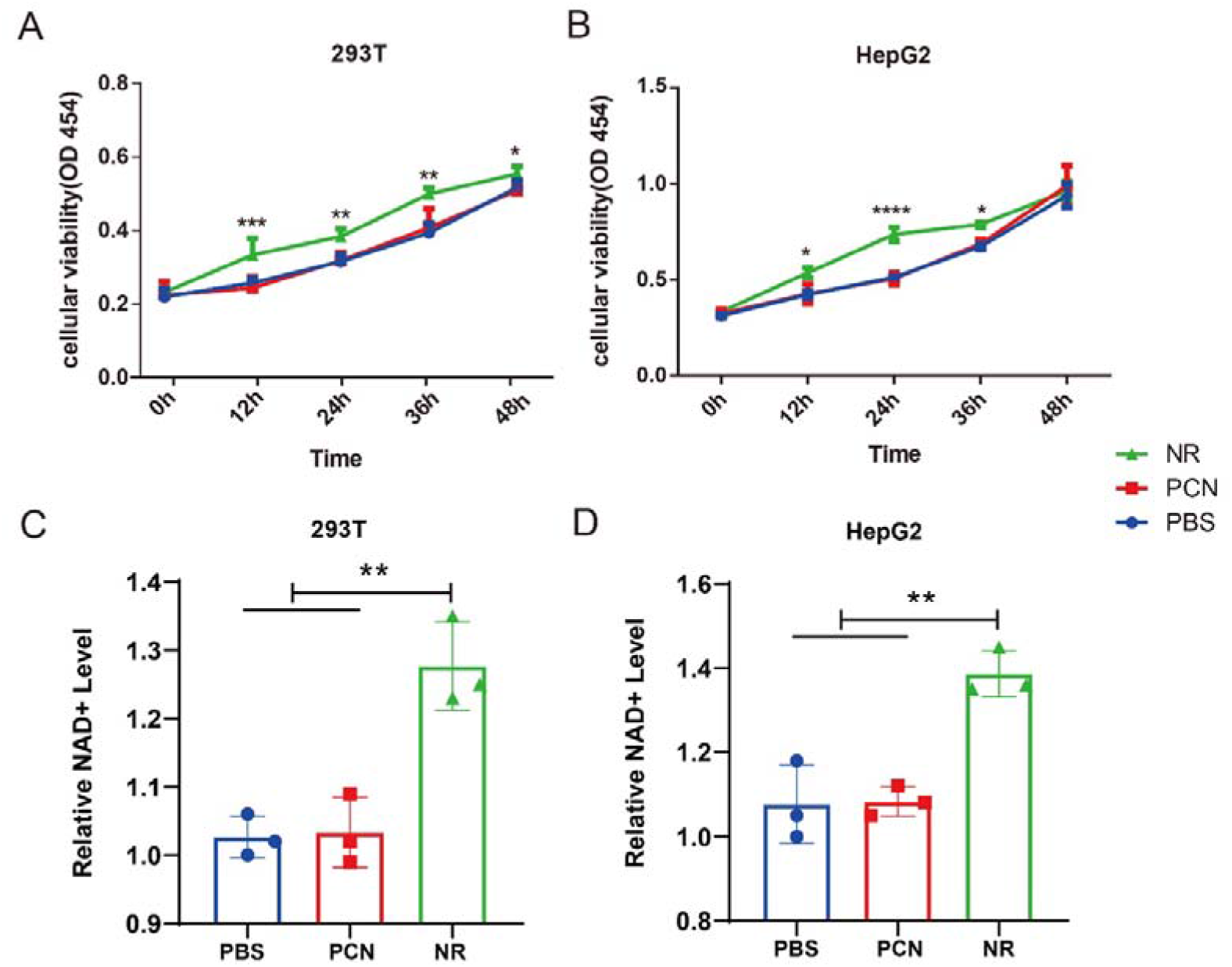
PCN does not affect the growth status and NAD^+^ level of 293T and HepG2 cells. A. Cell vitality of 293T cells after NR and PCN treatment. B. Cell vitality of HepG2 cells after NR and PCN treatment. C. NAD^+^ level in 293T cells after NR and PCN treatment. D. NAD^+^ level in HepG2 cells after NR and PCN treatment.

Because PncA in the intestinal flora plays an important role in the synthesis of NAD^+^ in the host intestine and liver, we next determined the effects of PCN treatment on the level of NAD^+^ in the liver and intestine of host mice. PCN reduced the utilization efficiency of NAM in the liver and intestine of the host, resulting in lower NAD^+^ levels (Fig. 2D, F). Given that PncA catalyzes the conversion of NAM to NA, we speculated that NAD^+^ synthesis using NA is more efficient than that using NAM in the liver and intestine. It is well known that NAM is present in sufficient amounts in the body as it is produced when NAD^+^ is consumed, and additional supplementation does not enhance NAD^+^ synthesis. Surprisingly, PCN also reduced the level of NAD^+^ in the hippocampus (Fig. 2F). According to RNA-Seq results, PCN affected the expression of several genes in the liver (Supplementary Fig. 2D) and significantly affected host immune processes, such as the intestinal immune network for IgA production and antigen processing (Supplementary Fig. 2E). To demonstrate that PCN regulates the NAD^+^ level in the host through its effect on the intestinal flora rather than directly affecting the host, 293T and HepG2 cells were treated with PCN and NR. The results showed that PCN had no effect on the growth and NAD^+^ level of these cells (Supplementary Fig. 3).

### Different *PncA* genotypes of *Escherichia coli* affect host liver NAD metabolism

Our previous experiments and the research ^11^ showed that the *PncA* gene in the intestinal flora plays an important role in the regulation of host NAD^+^. Therefore, we attempted to promote the synthesis of mammalian NAD^+^ using a new approach. To this end, we first supplemented gut bacteria that specifically depend on the deamidation NAD^+^ synthesis pathway. However, because controlling univariate variables for this experiment is challenging, we constructed induced PncA overexpression (PncA-OE) and PncA knockout (PncA-KO) transgenic *E. coli* strains. The PncA-KO strain was constructed by gene targeting technology (Supplementary Fig. 4A), and *PncA* gene KO from the E. coli genome (Supplementary Fig. 4B) was verified by PCR. Based on the gene expression of *PncA* in the three genotypes of *E. coli* (Supplementary Fig. 4C), *PncA* expression in the PncA-OE group was significantly higher than that in the other two groups.

**Supplementary Figure 4:**
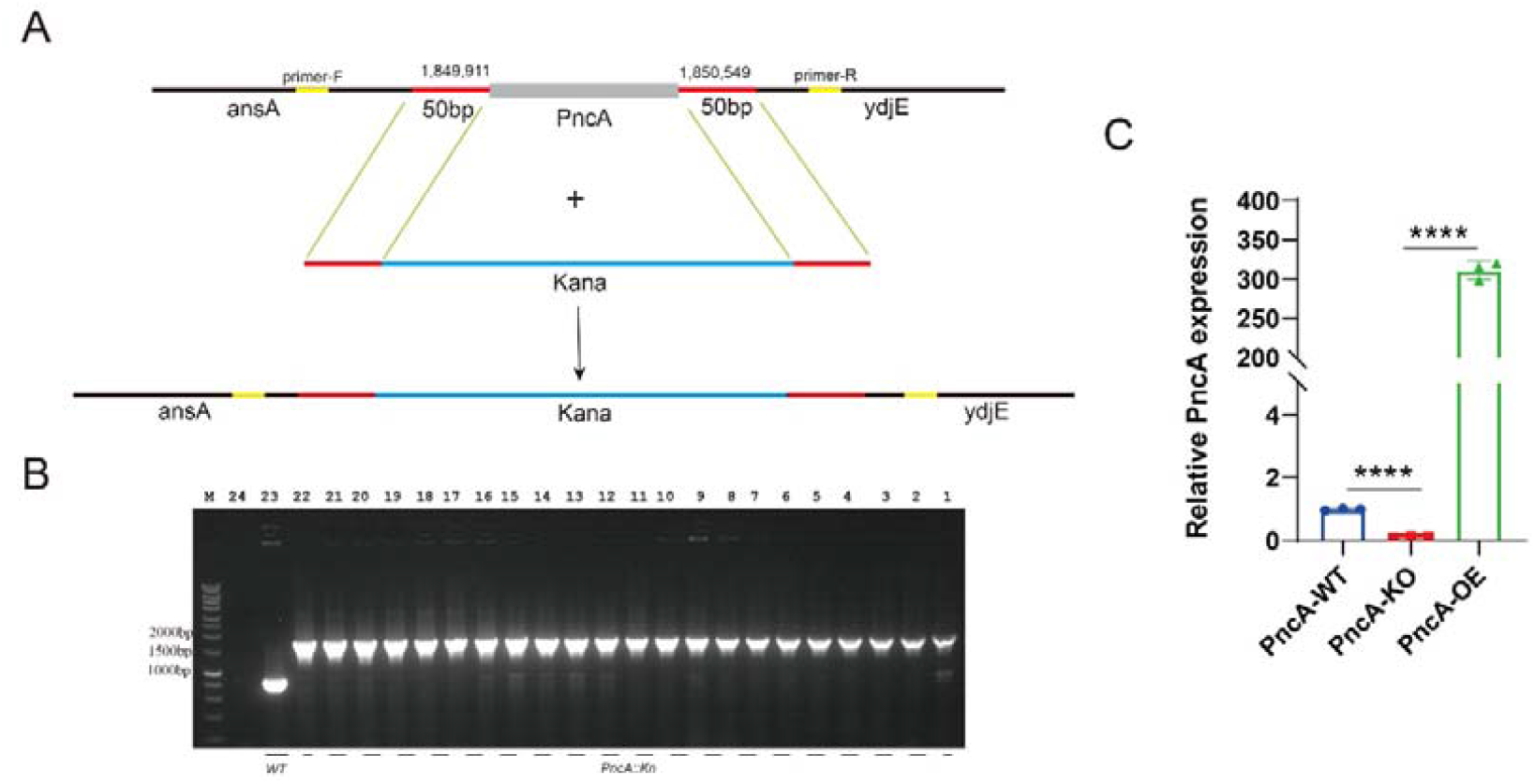
Construction and validation of *Escherichia coli* with different *PncA* genotypes. A. Diagram of *PncA* knockout *Escherichia coli* constructed by gene targeting technology. B. Verification of the knockout efficiency of *PncA* in *Escherichia coli*. C. Relative expression of *PncA* in wild type, PncA-KO, and PncA-OE *Escherichia coli*.

First, we performed an *in vitro* experiment. NAM was added to the culture medium of *E. coli*, and the bacteria were cultured for the indicated period. Bacteria were centrifuged, and the supernatant was collected for metabolome sequencing (Fig. 3A). As expected, PncA-OE *E. coli* released more NA in the bacterial culture medium than PncA-KO *E. coli*, and higher levels of NAD^+^ were also detected in the supernatant (Fig. 3B). To facilitate the colonization of exogenous *E. coli* in the intestines of mice *in vivo*, we first treated mice with cocktail antibiotics for 5 days to reduce endogenous bacteria. Next, mice were gavaged with PncA-OE and PncA-KO *E. coli* and divided into NAM-supplemented and non-NAM-supplemented groups (Fig. 3C). Previous studies showed that the efficiency of NA utilization for NAD^+^ synthesis in the liver is higher than that of NAM. Indeed, we found that the synthesis of NAD^+^ by PncA-OE *E. coli* was more efficient (Fig. 3D).

**Fig. 3:**
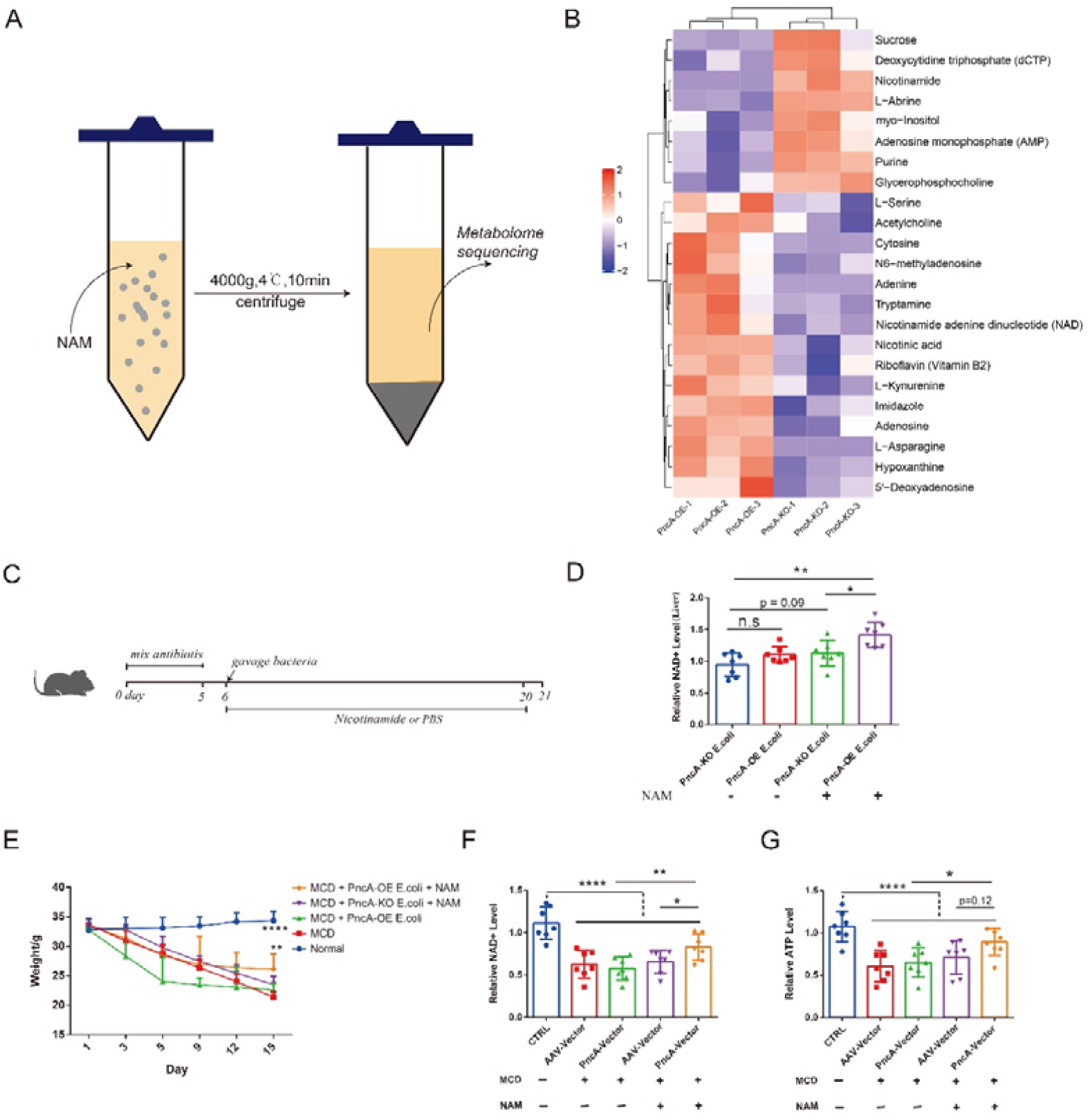
Different *PncA* genotypes of *Escherichia coli* affect host liver NAD metabolism. A. Schematic diagram of the *in vitro* experiment. B. The heatmap of the metabolites in the bacteria culture medium. C. Schematic diagram of the mouse experiment. D. Liver NAD^+^ levels after colonizing mice with different genotypes of *E. coli*. E. Body weight of MCD-induced non-alcoholic fatty liver model mice. F. Liver NAD^+^ level in non-alcoholic fatty liver model mice treated with different genotypes of bacteria. G. Relative content of ATP in mouse livers.

Enhancing the synthesis of NAD^+^ significantly improves NAFLD. Therefore, we further constructed NAFLD model mice using a methionine- and choline-deficient (MCD) diet to verify whether PncA-OE *E. coli* ameliorates this disease. The experimental results showed that MCD rapidly decreased the weight of mice (Fig. 3E) and significantly reduced liver NAD^+^ and ATP levels (Fig. 3F, G). In addition, after supplementation with PncA-OE *E. coli*, NAD^+^ and ATP levels in the liver were slightly increased compared with the control group (Fig. 3F, G).

### PncA overexpression in the liver by AAV improves hepatic lesions in mice

We found that supplementation with PncA-OE *E. coli* was not very effective in relieving NAFLD. However, we speculate that this may be due to the limited ability of bacteria to promote liver NAD^+^. Therefore, we optimized the sequence of *PncA* in *E. coli* for efficient expression in mammals. Then, a liver-specific AAV virus was used to carry the *PncA* gene into the liver for specific expression. We detected the gene expression of *PncA* in the mouse liver and found substantially high expression (Fig. 4A). Intriguingly, the NAD^+^ level in the PncA-OE group was significantly higher than that in the vector group and ∼5 times higher than that in the normal diet group (Fig. 4B). This result was particularly surprising because the widely used and highly efficient NAD^+^ precursors NR and NMN only increased the level of NAD^+^ in the liver by 1.5-to 2-fold. However, previous studies have shown that NA does not significantly enhance NAD^+^ levels in mouse livers. By comparing our experiment with previous experiments reported by others, we speculate that the effect of NA on NAD^+^ synthesis may be limited by the absorption efficiency of NA by liver cells. PncA expression in the liver promotes the transformation of NAM to NA in cells, maximizing the synthesis of NA to NAD^+^. This suggests that the deamidase PncA appears to have been abandoned during the evolution of the NAD^+^ synthesis pathway and has great potential for NAD^+^ synthesis in mammals.

**Fig. 4:**
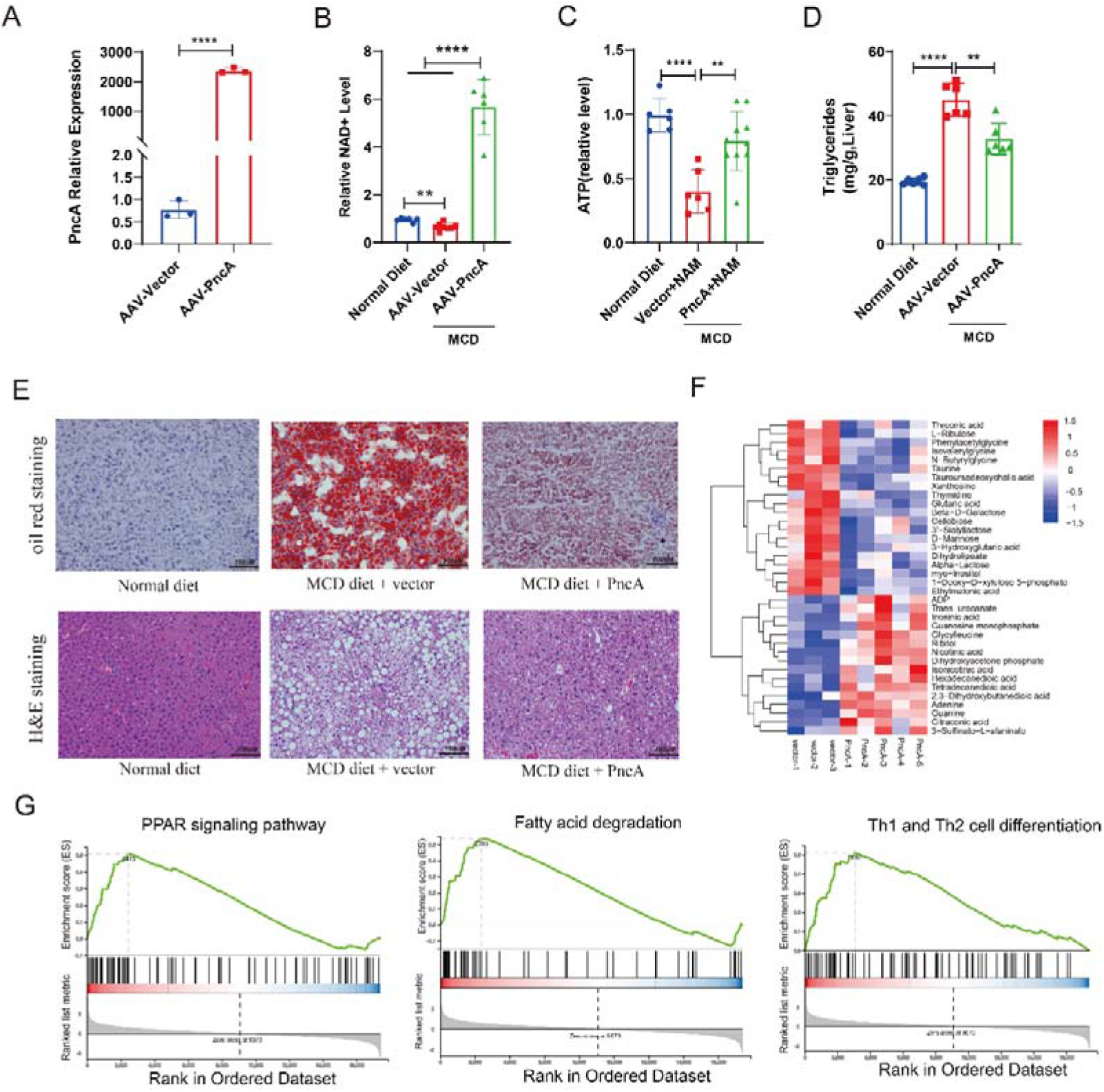
Improvement in mouse liver lesions by PncA overexpression via AAV. A. PncA expression in the livers of the control group and the PncA group. B. Relative content of NAD^+^ in mouse livers. C. Relative content of ATP in mouse livers. D. Content of triglyceride in the livers of mice. E. Representative Oil Red O staining (top) and hematoxylin and eosin (H&E) staining (bottom) of mouse liver sections. F. Heatmaps of metabolites with significant differences between the PncA and vector groups. G. GSEA analysis of the RNA-seq results.

In addition, compared with the vector group, mice expressing PncA showed a significantly increased ATP level and a significantly decreased triglyceride content after supplementation with the MCD diet (Fig. 4C, D). Furthermore, the results of Oil Red O staining and H&E staining showed that the overexpression of *PncA* in mouse livers greatly improved NAFLD-induced pathological changes (Fig. 4E). Based on the metabolome sequencing results, PCA revealed significant differences between the PncA and vector groups (Supplementary Fig. 5B), and the volcano diagram demonstrated a large number of differential metabolites between the two groups (Supplementary Fig. 5D). The NA amount in the liver of the PncA group was significantly increased (Fig. 4F), and the levels of other small molecules involved in nicotinate and NAM metabolism were also significantly altered (Supplementary Fig. 5C). Metabolites that differed between the two groups were mainly enriched in nicotinate and NAM metabolic pathways, followed by phenylalanine, tyrosine, and tryptophan biosynthesis and other pathways related to lipid metabolism (Supplementary Fig. 5E). RNA-seq analysis results showed that the expression of some genes involved in lipid metabolism was significantly changed (Supplementary Fig. 5A), and gene set enrichment analysis revealed that PncA increased the expression of genes mainly involved the PPAR signaling pathway, fatty acid degradation, and other pathways that promote fat metabolism (Fig. 4G). To our surprise, the highly expressed genes in the PncA group were also significantly enriched in the Th1 and Th2 cell differentiation pathways, indicating that *PncA* genes may play an important role in T cell differentiation.

**Supplementary Figure 5:**
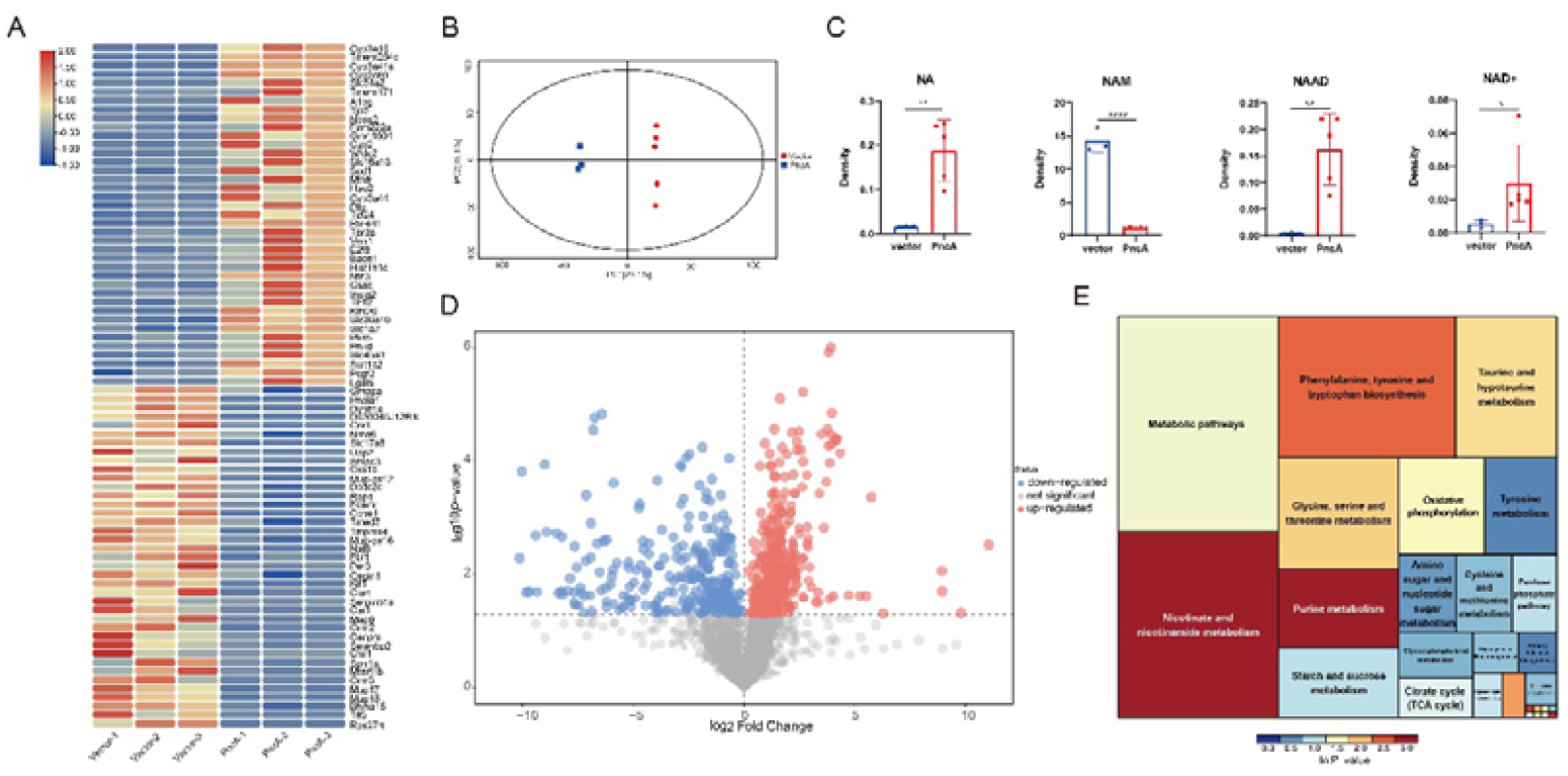
PncA affected the genes and metabolites related to lipid metabolism in NAFLD model mice. A. Heatmap of differently expressed genes between PncA and vector groups. B. PCA of metabolites in PncA and vector groups. C. Relative contents of NA, NAM, NAAD, and NAD^+^ in mouse livers. D. Volcano plot of metabolites in PncA and vector groups. The colored dots represent differentially regulated metabolites between the two groups. E. Enriched KEGG pathways of differentially regulated metabolites.

To verify that direct NA supplementation in mice could not reappear the effect by expressing PncA in the liver, we treated mice with the MCD diet and NA. The weight of mice decreased throughout the experiment (Supplementary Fig. 6A), and NA had no obvious effect on the NAD^+^ level in the liver of mice (Supplementary Fig. 6B). Therefore, direct NA supplementation does not increase the level of NAD^+^.

**Supplementary Figure 6:**
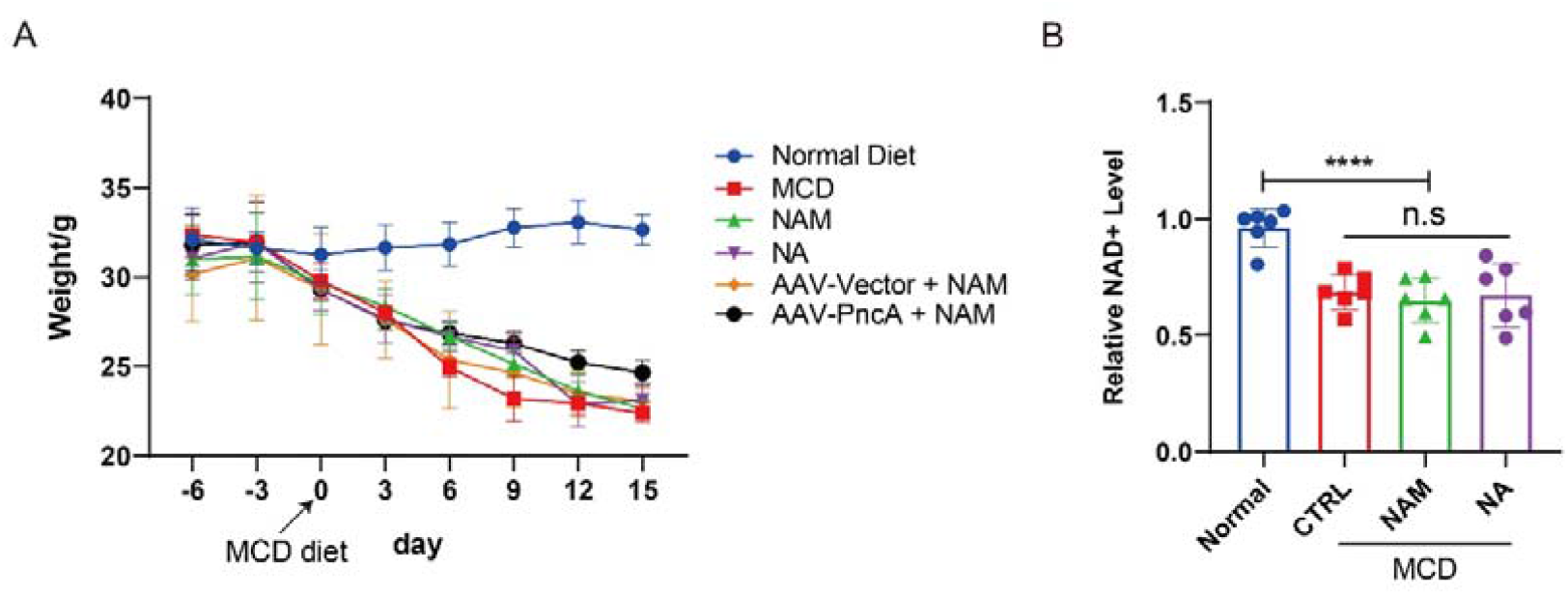
NA did not increase the NAD+ level in NAFLD mice. A. Body weight of mice during the experiment after supplementation with the MCD diet. The weight of mice gradually decreased. B. Relative NAD+ level in the liver in the normal group and MCD group treated with PBS, NAM, and NA.

## Discussion

Coenzyme NAD^+^ plays a key role in cellular biology and adaptive stress responses. Its depletion is a basic feature of aging and may lead to various chronic diseases ^20^. NAD^+^ supplementation alleviates several aging-related diseases and even prolongs the lifespan of mice ^21^. Maintaining NAD^+^ levels is essential for the function of high-energy-demanding cells and mature neurons ^22^. NAD^+^ levels are substantially decreased in major neurodegenerative diseases, such as Alzheimer’s disease, Parkinson’s disease, cardiovascular disease, and muscle atrophy ^23^. Increasing evidence has shown that NAD^+^ is significantly reduced in various tissues during aging, and determining how to efficiently increase cellular NAD^+^ levels through physiological and pharmacological methods and prevent age-related diseases has become a research topic of high interest in recent years. Similarly, identifying new and efficient methods to promote NAD^+^ synthesis is vital.

Currently, the most efficient NAD^+^ synthesis pathway in mammals is a controversial issue. In addition, different organs depend on different NAD^+^ precursors ^24^. Studies have reported that the liver and kidney use three NAD^+^ synthesis pathways, the spleen, small intestine, and pancreas mainly rely on NA and NAM, and the heart, lung, brain, muscle, and white adipose tissue primarily use the NAM pathway ^25^. Regarding the most efficient precursor, some articles have reported that the efficiency of NAD^+^ synthesis with NA is higher than that with NAM, but there is still no clear conclusion ^26^. However, the most widely used NAD^+^ precursors are NR and NMN. After these two precursors enter the cell, NAD^+^ is synthesized through one or two enzymatic reactions, and it avoids being affected by rate-limiting enzymes, such as NAMPT. These two precursors have been used for various interventions, such as alleviating neurodegenerative diseases, improving hearing, treating diabetes and NAFLD, delaying aging, and extending lifespans ^27-31^.

Cells have different absorption efficiencies for various NAD^+^ precursors. NAM directly enters the cell through free diffusion, whereas NA, NR, NMN, and other precursors require the assistance of membrane proteins, resulting in a lower absorption efficiency compared with NAM ^32^. However, NAM is believed to be present in sufficient amounts in the body because NAD^+^ is decomposed to produce NAM, which re-enters the NAD^+^ synthesis pathway. In addition, NAM is an inhibitor of the SIRT family. Excessive concentrations of NAM inhibit the activity of sir2 and shorten the lifespan of *Saccharomyces cerevisiae* ^33^. In contrast, studies have shown that NAM supplementation does not increase the lifespan of mice ^34^. However, PncA overexpression in *Drosophila* protects neurons and extends their lifespan ^35^. PncA also increases the lifespan of *Caenorhabditis elegans* ^36^. In this study, we used an AAV to express PncA in the liver, an organ that relies on multiple NAD^+^ precursors. In this way, we replenished the liver with the deamidase that was discarded during evolution. The results showed that the deamidase was highly active in mammalian cells, and this approach increased the level of NAD^+^ to a greater extent than NR or NMN.

Furthermore, the efficiency of using NA for NAD^+^ synthesis appears to be the highest. In addition, PncA processes excessive NAM in the liver, thereby releasing the inhibition of SIRT1 by NAM and increasing the activity of SIRT1. Based on the above results, we speculate that PncA will also substantially increase the NAD^+^ levels in other organs that rely on NA to synthesize NAD^+^. This prediction needs to be further studied. Moreover, we demonstrated that PncA significantly improves NAFLD in mice, indicating that PncA provides a promising potential target for the treatment of various diseases related to NAD^+^ deficiency.

## Methods

### Reagents

Nicotinamide (#HY-B0150), nicotinic acid (#HY-B0143), and nicotinamide riboside (#HY-123033) were obtained from MedChemExpress (Monmouth Junction, NJ, USA). Pyrazinecarbonitrile (#243-369-5) was obtained from Sigma-Aldrich (St. Louis, MO, USA), and the methionine-choline deficient (MCD) diet (#TD.90262) was obtained from Harlan Teklad. Brain heart infusion (#HB8297-5), MRS (#HB0384-1), and TPY (#HB8570) were obtained from Hopebio-Technology (Qingdao, China).

### Cell lines

293T female embryonic kidney cells and HepG2 liver cancer cells were obtained from ATCC and cultured in Dulbecco’ Modified Eagle’s Medium (Thermo Fisher Scientific, Waltham, MA, USA) supplemented with 10% fetal bovine serum (FBS) and 100 mg/ml penicillin/streptomycin. Each cell line was maintained in a 5% CO_2_ atmosphere at 37°C. The mycoplasma contamination status of all cultures was monitored monthly by PCR.

### Bacteria strains

*A. muciniphila* (ATCC BAA-835) and *Lactobacillus salivarius* (ATCC 1174) were cultured anaerobically in brain heart infusion and MRS medium, respectively. *Bifidobacterium longum* (#1.2186) was obtained from the China General Microbiological Culture Collection Center (CGMCC) (Beijing, China) and cultured anaerobically in TPY medium.

### Mice

WT C57BL/6J mice were purchased from Cyagen Biosciences Inc. Mice were maintained under a 12-h light/dark cycle and fed a standard chow diet in the specific pathogen-free (SPF) facility at the Laboratory Animal Research Center, Tongji University. For bacterial and AAV infection, 8-to 10-week-old female C57BL/6J mice were used. All mouse experiments were carried out following the national guidelines for the housing and care of laboratory animals (Ministry of Health, China), and the protocol complied with institutional regulations after review and approval by the Institutional Animal Ethics Committee at the Laboratory Animal Research Center, Tongji University (TJAA09220102).

### Mice infection

Eight-week-old C57BL/6 female mice were gavaged with either regular water or autoclaved water containing an antibiotic cocktail (1 g/L ampicillin, 1 g/L neomycin, 1 g/L metronidazole, 500 mg/L vancomycin) for five days and then given regular water. *E. coli* strains were cultured as described above and quantified prior to infection. For bacterial colonization assays, the mice were infected intragastrically with 1×10^9^ PncA-OE or PncA-KO *E. coli* strains (in 0.2 mL PBS) every 3 days until the end of the experiment. For the group with NAD^+^ precursors, NAM (4 mg/kg/day) and NA 4 mg/kg/day were delivered via gavage. AAV (1 × 10^12^ pfu/ml adenovirus in 100 μL PBS) expressing *PncA* (AAV-PncA) or vector (AAV-vector) were injected into mice via their tail vein. Animals were sacrificed 60 days after injection. The liver was removed and stored at −80°C until use.

### Cell Transfection

Transient transfection was performed using LipoFiter 3.0 (Hanbio Biotechnology Co., Ltd) following the manufacturer’s instructions.

### Preparation of AAV

293T cells were used to package adenoviruses using a three-plasmid system, including pAAV-RC, pHelper, and shuttle plasmids (with or without the target gene). 293T cells were subcultured into 100-mm plates for transfection. Transfection was performed when the cell density reached 80%–90% with the following transfection complex reagents: pAAV-RC 10 µg, pHelper 20 µg, shuttle plasmid 10 µg, and Lipofiter™ (HB-TRCF-1000, Hanbio Biotechnology) 120 µL. Fresh complete medium containing 10% FBS was replaced 6 h after transfection. Seventy-two hours after transfection, cells containing AAV particles were gently removed using a cell scraper, collected into a 15-mL centrifuge tube, and centrifuged at 150 g for 3 min. Cells were collected, and the culture supernatant was removed. Cells were washed with PBS once and re-suspended with 300 μL PBS. The cells were frozen and thawed in liquid nitrogen and 37°C three times and then centrifuged at 2000 g for 5 min at 4°C to remove cell debris. The lysis supernatant containing AAV particles was collected, and 0.1 μL Benonase (9025-65-4, Merck) was added to each 1 mL of crude virus extract to remove cell genome and plasmid DNA. The cells were centrifuged at 600 g for 10 min at 4°C, and the supernatant was collected for column purification (V1469-01, Biomiga). The 4-mL AAV liquid samples purified by the column were added to the ultrafiltration tube and centrifuged at 1400 g for 30 min to obtain approximately 1 mL of purified AAV, which was stored at −80°C until use.

### Untargeted metabolomics and LC-MS/MS

For LC-MS/MS, 50-mg samples were weighed and placed in an EP tube, and 1000 μL extract solution (methanol:acetonitrile:water = 2:2:1, with an isotopically-labeled internal standard mixture) were added. Then, the samples were homogenized at 35 Hz for 4 min and sonicated for 5 min in an ice-water bath. The homogenization and sonication cycles were repeated three times. Then, the samples were incubated for 1 h at −40°C and centrifuged at 12000 rpm for 15 min at 4°C. The resulting supernatant was transferred to a fresh glass vial for analysis.

LC-MS/MS analyses were performed using a UHPLC system (Vanquish, Thermo Fisher Scientific) with a UPLC BEH Amide column (2.1 mm × 100 mm, 1.7 μm) coupled to a Q Exactive HFX mass spectrometer (Orbitrap MS, Thermo Fisher Scientific). The mobile phase consisted of 25 mmol/L ammonium acetate and 25 mmol/L ammonia hydroxide in water (pH = 9.75) (A) and acetonitrile (B). The auto-sampler temperature was 4°C, and the injection volume was 2 μL. The QE HFX mass spectrometer was applied to acquire MS/MS spectra using the information-dependent acquisition (IDA) mode with acquisition software (Xcalibur, Thermo Fisher Scientific). In this mode, the acquisition software continuously evaluates the full MS spectrum. The ESI source conditions were set as follows: sheath gas flow rate = 30 Arb, Aux gas flow rate = 25 Arb, capillary temperature = 350°C, full MS resolution = 60000, MS/MS resolution = 7500, collision energy = 10/30/60 in the NCE mode, and spray voltage = 3.6 kV (positive) or −3.2 kV (negative).

The raw data were converted to the mzXML format using ProteoWizard and processed with an in-house program (developed using R and based on XCMS) for peak detection, extraction, alignment, and integration. Then, an in-house MS2 database (BiotreeDB) was applied for metabolite annotation. The cutoff for annotation was set at 0.3.

### RNA isolation, RNA-seq, and data processing

Total RNA was extracted from tissues using Trizol (Invitrogen, Carlsbad, CA, USA) in accordance with the manual’s instructions. Oligo (dT)-attached magnetic beads were used to purify mRNA. Purified mRNA was fragmented into small pieces with fragment buffer at the appropriate temperature. Then, first-strand cDNA was generated using random hexamer-primed reverse transcription, followed by second-strand cDNA synthesis. Afterwards, A-Tailing Mix and RNA Index Adapters were added by incubating to end repair. The cDNA fragments obtained in the previous step were amplified by PCR, and products were purified using Ampure XP Beads and then dissolved in EB solution. The product was validated on the Agilent Technologies 2100 bioanalyzer for quality control. The double-stranded PCR products from the previous step were heated, denatured, and circularized by the splint oligo sequence to obtain the final library. Single-strand circle DNA (ssCir DNA) was formatted as the final library. The final library was amplified with phi29 to generate DNA nanoballs (DNBs) containing more than 300 copies of one molecule. DNBs were loaded into the patterned nanoarray, and paired-end 50 base pair reads were generated on the BGIseq500 platform (BGI, Shenzhen, China).

The sequencing data were filtered with SOAPnuke (v1.5.2) ^37^ by (1) removing reads containing sequencing adapters, (2) removing reads with a low-quality base ratio (base quality less than or equal to 5) more than 20%, and (3) removing reads with an unknown base (N’ base) ratio more than 5%. Afterward, clean reads were obtained and stored in FASTQ format. The clean reads were mapped to the reference genome using HISAT2 (v2.0.4) ^38^. Bowtie2 (v2.2.5) ^39^ was applied to align the clean reads to the reference coding gene set, and then the gene expression level was calculated by RSEM (v1.2.12) ^40^. The heatmap was drawn by pheatmap (v1.0.8) according to the gene expression in different samples. Differential expression analysis was performed using the DESeq2(v1.4.5) ^41^ with a Q value < 0.05. To gain insight into phenotypic changes, GO (http://www.geneontology.org/) and KEGG (https://www.kegg.jp/) enrichment analysis of annotated differentially expressed genes were performed by Phyper (https://en.wikipedia.org/wiki/Hypergeometric distribution) based on the Hypergeometric test. The significant levels of terms and pathways were corrected using the Q value with a rigorous threshold (Q value < 0.05) by the Bonferroni test.

### qRT-PCR

RNA was treated with DNase, and 1 μg RNA was used for reverse transcription. cDNA diluted 10× was used for reverse transcription-quantitative PCR (RT-qPCR) reactions. RT-qPCR reactions were performed using the Light-Cycler system (Roche Diagnostics GmbH, Rotkreuz, Switzerland) and a qPCR Supermix (Vzayme, Nanjing, China) with the indicated primers. An average of at least three technical repeats was used for each biological data point. The *ropB* gene was used as the reference gene, and the following primers were used: *ropB*-F, CTGCGCGAAGAAATCGAAGG, *ropB*-R: TTTCGCCAACGGAACGGATA and *PncA*-F: TGATCGCCAGCCAAGACT, *PncA*-R: AGCATCCAGCACC GTGAA.

### 16S rRNA sequencing and analysis

#### Genomic DNA extraction

Microbial community DNA was extracted using a MagPure Stool DNA KF kit B (Magen, Guangzhou, China) following the manufacturer’s instructions. DNA was quantified with a Qubit Fluorometer using a Qubit dsDNA BR Assay kit (Invitrogen, Carlsbad, CA, USA), and the quality was assessed by running an aliquot on a 1% agarose gel.

#### Library Construction

Variable regions V3–V4 of the bacterial 16S rRNA gene were amplified with the degenerate PCR primers 341F (5’-ACTCCTACGGGAGGCAGCAG-3’) and 806R (5’-GGACTACHVGGGTWTCTAAT-3’). Both forward and reverse primers were tagged with Illumina adapter, pad, and linker sequences. PCR enrichment was performed in a 50-µL reaction containing 30 ng template, fusion PCR primer, and PCR master mix. PCR cycling conditions were as follows: 94°C for 3 min, 30 cycles of 94°C for 30 seconds, 56°C for 45 seconds, and 72°C for 45 seconds, and a final extension for 10 min at 72°C. The PCR products were purified with AmpureXP beads and eluted in Elution buffer. Libraries were qualified with the Agilent 2100 bioanalyzer (Agilent, Santa Clara, CA, United States). The validated libraries were used for sequencing on an Illumina MiSeq platform (BGI, Shenzhen, China) following the standard pipelines of Illumina, and 2 × 300 bp paired-end reads were generated.

#### Oil Red staining and H&E staining

Hepatic tissue was cut into small pieces, fixed in 4% paraformaldehyde for 4 h, and embedded in OCT (Leica Camera AG, Wetzlar, Germany). Frozen sections (4 µm thickness) were made using a cryostat, and the samples were fixed with 4% paraformaldehyde for an additional 30 min. The slides were washed in distilled water and stained with Oil Red O for 15 min. Next, the slides were counterstained with hematoxylin for 10 s to identify the nuclei. For H&E staining, the slides were first stained in hematoxylin for 3–8 min and then counterstained with eosin for 1–3 min. The histological images were acquired with a light microscope (Olympus; Tokyo, Japan).

#### NAD^+^ detection

NAD^+^ detection was carried out using the EnzyChrom TM NAD^+^/NADH^+^ Assay Kit (#E2ND-100; BioAssay Systems, Hayward, CA, USA) following the standard steps provided. Protein concentration was used to normalize the NAD^+^ content.

#### ATP detection

For the measurement of ATP levels, 100-mg liver samples were lysed in 1 ml lysis buffer provided by the ATP Assay Kit (#S0026; Beyotime, Jiangsu, China). Liver ATP levels were evaluated by luciferase activity, as shown in the standard protocol provided by the ATP Assay Kit.

#### Triglyceride detection

Liver triglycerides were assayed using a triglyceride assay kit (#E1025; Applygen Technologies, Beijing, China).

#### Statistical analysis

All data were analyzed using appropriate statistical methods with GraphPad Prism 8. Differences between two groups were evaluated using Student’s t-test. A P value of less than 0.05 was considered statistically significant. All data are presented as the mean ± SD. Sample sizes were selected without performing statistical tests. No data were excluded when conducting the final statistical analysis. *P < 0.05, **P ≤ 0.01, and ***P ≤ 0.001 unless stated otherwise.

## Data availability statement

The RNA-seq raw sequencing data files generated in this study are available in the NCBI’s Sequence Read Archive (SRA) under BioProject accession number PRJNA780659. The original 16S rRNA sequence data are available at the NCBI by accession number PRJNA780391.

## Acknowledgments

This work was supported by grants from the National Key Research and Development Program (2020YFC2002800), the Fundamental Research Funds for the Central University (22120210584), and Major Program of Development Fund for Shanghai Zhangjiang National Innovation Demonstration Zone<Stem Cell Strategic Biobank and Stem Cell Clinical Technology Transformation Platform> (ZJ2018-ZD-004).

## Author information

## Contributions

L.H.L and F.S.Y designed the study; F.S.Y. and G.L.L analyzed data; L.H.L supervised the entire project; and all authors approved the final version of the manuscript.

## Corresponding authors

Correspondence to Hailiang Liu.

